# HRMAS ^13^C NMR and genome-scale metabolic modeling identify threonine as a preferred dual redox substrate for *Clostridioides difficile*

**DOI:** 10.1101/2023.09.18.558167

**Authors:** Aidan Pavao, Ella Zhang, Auriane Monestier, Johann Peltier, Bruno Dupuy, Leo Cheng, Lynn Bry

## Abstract

Stickland-fermenting *Clostridia* preferentially ferment amino acids to generate energy and anabolic substrates for growth. In gut ecosystems, these species prefer dual redox substrates, particularly mucin-abundant leucine. Here, we establish how theronine, a more prevalent, mucin-abundant substrate, supports dual redox metabolism in the pathogen *Clostridioides difficile*. Real-time, High-Resolution Magic Angle Spinning NMR spectroscopy, with dynamic flux balance analyses, inferred dynamic recruitment of four distinct threonine fermentation pathways, including ones with intermediate accrual that supported changing cellular needs for energy, redox metabolism, nitrogen cycling, and growth. Model predictions with ^13^C isotopomer analyses of [U-^13^C]threonine metabolites inferred threonine’s reduction to butyrate through the reductive leucine pathway, a finding confirmed by deletion of the *hadA* 2-hydroxyisocaproate CoA transferase. *In vivo* metabolomic and metatranscriptomic analyses illustrate how threonine metabolism in *C. difficile* and the protective commensal *Paraclostridium bifermentans* impacts pathogen colonization and growth, expanding the range of dual-redox substrates that modulate host risks for disease.

## Introduction

Anaerobic and sporulating *Clostridia* occupy diverse metabolic niches of clinical, environmental, and industrial importance[1-3]. The pathogen *Clostridioides difficile* leverages its complex metabolism to harness substrates of dietary, host, and microbiota origin for colonization and rapid growth, particularly in enriched gut nutrient environments that occur after antibiotic ablation of the microbiota [4-7].

Thought to have evolved in the earth’s early amino acid-rich ecosystems [4, 8-10], the Stickland pathways oxidize preferred substrates including branched-chain, aromatic, and pyruvogenic amino acids to generate ATP, proton gradients, and anabolic substrates, while reducing amino acids such as leucine, proline, and glycine through defined reductase enzymes that extract further energy for cellular needs [4]. The Stickland half reactions enhance the pathogen’s metabolic flexibility by recruiting distinct redox reactions to support high-flux glycolytic, mixed acid, or other forms of fermentation. This metabolic flexibility reflects *C. difficile’s* preference for dual-redox amino acids such as leucine which has dedicated oxidative and reductive pathways to support cellular needs [8].

Notably, *C. difficile’s* preferred reductive substrates leucine, proline, and glycine are prevalent in gut mucins[11]. However, threonine is the most abundant, gut-available amino acid, occurring as a free amino acid as well as in mucin backbones, where it comprises up to 30% of mucin amino acid content[11-13]. *C. difficile* actively consumes threonine *in vitro* and *in vivo* [14, 15]. Prior studies of *C. difficile’s* threonine metabolism have shown reduction to 2-oxobutyrate via threonine dehydratase, oxidative metabolism to propionate and CO_2_, and reduction to 2-hydroxybutyrate [14, 16]. In other *Clostridia*, including *Clostridium propionicum*, the reductive lactate-acrylate-propionate (LAP) pathway metabolizes threonine to butyrate [17-19]. While LAP activity has not been observed in *C. difficile* [20], studies by Eldsen *et al*. demonstrated 2-3 fold increases in butyrate with threonine supplementation, suggesting an unknown reductive pathway[21].

Using whole-cell and real-time High-Resolution Magic Angle Spinning (HRMAS) Nuclear Magnetic Resonance (NMR) spectroscopy with uniformly carbon-13 labeled [U-^13^C]threonine[20], we identify dynamic recruitment of four distinct oxidative and reductive pathways. including ones that accrue and then consume intermediates to support changing cellular needs in energy generation, redox balance, nitrogen cycling, and growth. Threonine’s pathways further support redox metabolism during a critical gap between high flux Stickland metabolism of branched chain amino acids and proline, and prior to the ramp-up of glycolytic metabolism. Model predictions inferred reductive threonine metabolism to butyrate by the 2-hydroxyisocaproyl CoA-transferase enzyme of the reductive leucine pathway, a finding confirmed in an isogenic *ΔhadA* strain. *In vivo*, we illustrate functions of threonine metabolism in *C. difficile* colonization, progression to disease, and in molecular approaches for disease prevention with the protective Stickland-fermenting commensal *Paraclostridium bifermentans*[22].

## Results

### *C. difficile’s* threonine metabolism supports sequential functions in nitrogen handling, ATP production, and redox metabolism

Real-time HRMAS ^13^C-NMR of *C. difficile* grown in Modified Minimal Media (MMM) with 15mM [U-^13^C]threonine revealed rapid consumption of threonine between 3-16 hours of growth (Fig. 1a,b). Analyses detected [^13^C]2-aminobutyrate by 6 hours, peak signal over 12-15 hours, with subsequent decrease into stationary phase (Fig. 1a,b; Fig. 2a). *C. difficile’s* oxidation of [U-^13^C]threonine to [U-^13^C]propionate and ^13^CO_2_ occurred over 6-16 hours, followed by [U-^13^C]propionate’s reduction to [U-^13^C]*n-*propanol over 12-36 hours (Fig. 1a,b; Fig. 2b,c). Reductive products [^13^C]2-hydroxybutyrate and [^13^C]butyrate arose at 11 hours through 36 hours (Fig. 1a,b; Fig. 2d), followed by [^13^C]ethanol and [^13^C]glycine at 14 and 16 hours, respectively (Fig. 1a; Fig. 2e).

**Figure 1.**
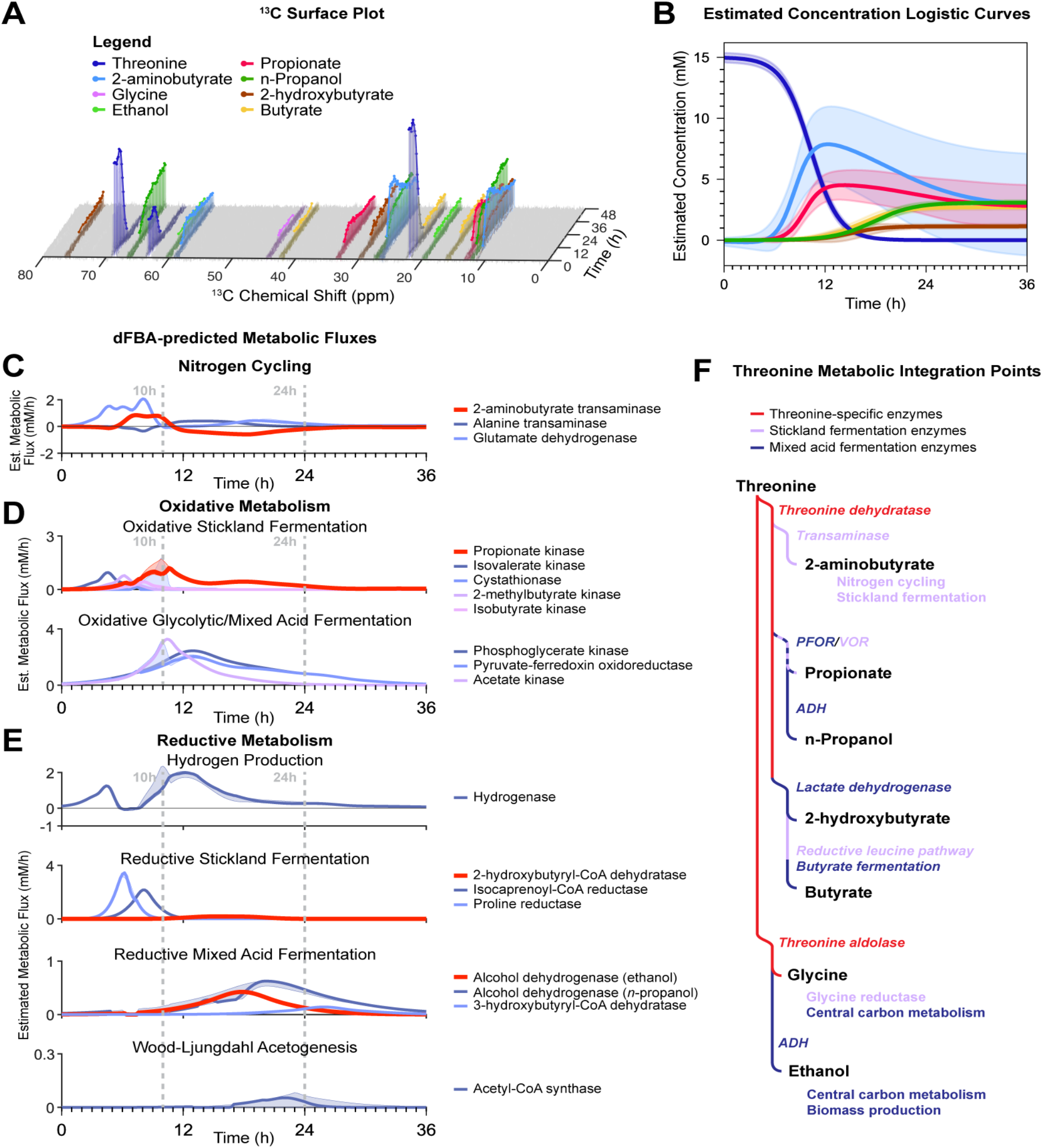
HRMAS ^13^C-NMR and dFBA resolve real-time fermentation of [U-^13^C]threonine by *C. difficile*. (**A**) Representative ^13^C surface plot of [U-^13^C]threonine metabolism. X-axis: ^13^C chemical shift; Y-axis: time in hours, Z-axis: NMR signal. (**B**) Logistic curves depicting estimated concentration of [U-^13^C]threonine and its products over time. Shaded regions depict 95% confidence interval across three experimental replicates; butyrate and 2-hydroxybutyrate curves show estimated curves averaged across replicates (n=2) with signal to support curve estimation and ones (n=1) with detectable metabolite signal that fell below thresholds for logistic curve fitting. (**C-E**) dFBA-predicted metabolic fluxes for key metabolic and cellular functions. Red trajectories indicate estimated fluxes inferred directly from [U-^13^C]threonine metabolite signals. Reactions shown in (**C**) show nitrogen cycling, (**D**) oxidative metabolism, and (**E**) reductive metabolism; X-axis: time in hours, Y-axis: estimated metabolic flux in mM/h. (**F**) Routemap of threonine metabolism indicating metabolic integration points among cellular systems. Italic text indicates enzymes associated with biochemical pathway branch points and associated cellular systems; colored labels indicate enzyme category as shown in the key at the top.

**Figure 2.**
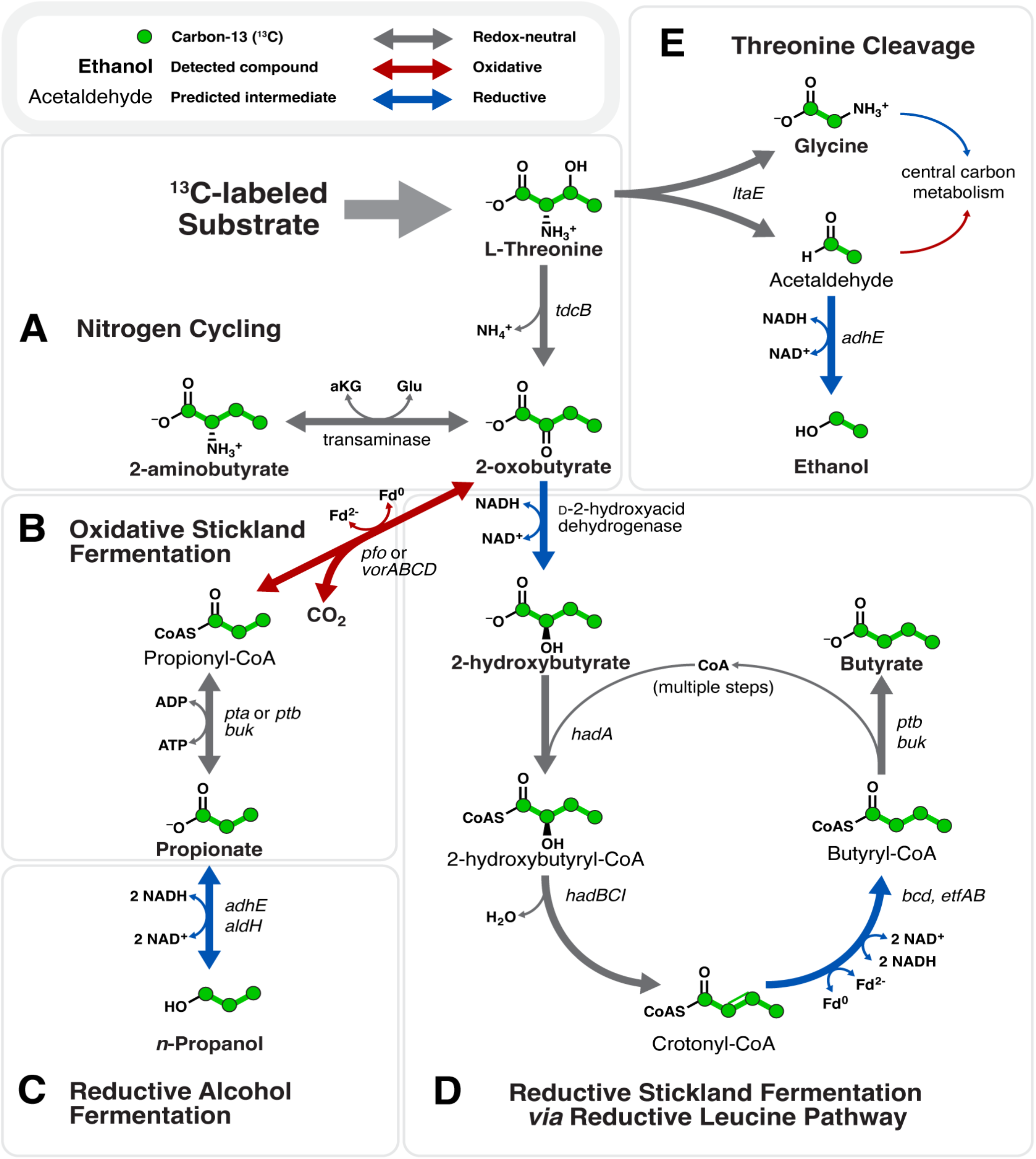
Threonine fermentation pathways in *C. difficile*. Enzyme encoding genes are shown to the right of, or below, reaction arrows. (**A**) Redox-neutral nitrogen cycling of threonine’s deamination to 2-oxobutyrate, and subsequent capacity for 2-oxobutyrate to accept amino nitrogen groups to produce 2-aminobutyrate. (**B**) Oxidative threonine fermentation to propionate followed by (**C**) reduction to *n*-propanol. (**D**) Proposed reduction of threonine to 2-hydroxybutyrate and butyrate via the reductive leucine pathway and the isocaproyl-CoA transferase encoded by *hadA*. Threonine cleavage to glycine and acetaldehyde via threonine aldolase, with subsequent reduction of acetaldehyde to ethanol.

Dynamic Flux Balance Analysis (dFBA) using NMR-informed constraints from [U-^13^C]threonine metabolism inferred changing support among cellular functions over phases of growth. dFBA simulations predicted initial threonine metabolism to 2-aminobutyrate, supporting nitrogen handling to sustain co-occurring Stickland fermentations, and providing an additional energy-efficient pathway to enhance alanine production from glycolytic-origin pyruvate (Fig. 1c), enabling pathogen growth during co-occurring Stickland and glycolytic metabolism[20].

Threonine’s oxidation to propionate over 8-12 hours sustained oxidative Stickland metabolism during a gap between the depletion of preferred branched-chain amino acid substrates, while glycolytic metabolism came online (Fig. 1d). Over 10-24 hours, threonine fermentation again switched to support reductive metabolism, producing butyrate from 2-aminobutyrate, and *n-* propanol from propionate, regenerating electron carriers as reductive flux from carbohydrate fermentations, producing lactate from pyruvate, and ethanol and butyrate from acetate, ramped up (Fig. 1e). Metabolism in this phase illustrated threonine’s capacity to cover gaps in oxidative and reductive metabolism as *C. difficile* switched among other amino acid and carbohydrate substrates.

Multiplet analyses of [^13^C]butyrate isotopologues infer activity of the reductive leucine pathway To identify pathway(s) responsible for threonine reduction to butyrate, labeling patterns of [^13^C]butyrate were evaluated to determine whether the carbons originated solely from threonine’s U-^13^C carbon backbone, or contained ^12^C contributions from glucose, glycine, and other natural abundance sources. Multiplet analysis of butyrate’s ^13^C isotopologues revealed ^13^C carbons at positions C2, C3, and C4, and a 1:1.4 ratio of ^12^C to ^13^C at C1, indicating production of [2,3,4-^13^C]butyrate and [U-^13^C]butyrate (Fig. 3a-d, black spectra). Analyses of ^13^C-labeleing in the upstream substrate 2-hydroxybutyrate (Fig. 3e-h, black spectra) showed [2,3,4-^13^C]hydroxybutyrate and [U-^13^C]hydroxybutyrate in a 1:1.2 ratio, indicating 2-hydroxybutyrate’s direct metabolism to butyrate without carbon rearrangement. Predictions inferred flux through the reductive leucine pathway to produce the resulting isotopologues of 2-hydroxybutyrate and butyrate from threonine.

**Figure 3.**
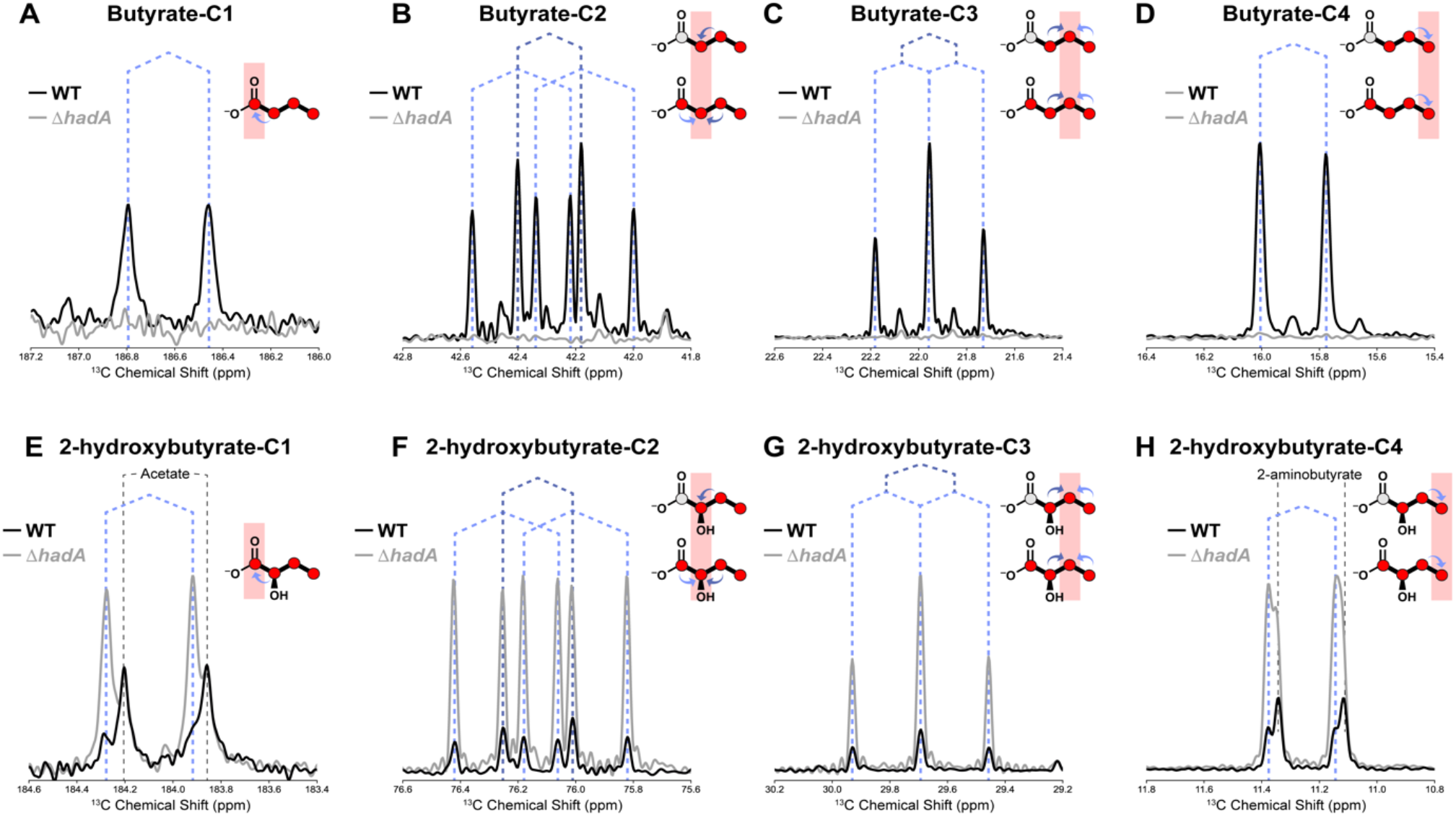
HRMAS ^13^C-NMR multiplet analysis confirms threonine’s fermentation to butyrate via metabolism of 2-hydroxybutyrate in the reductive leucine pathway. ^13^C-NMR signal of butyrate carbons in supernatants of *C. difficile* ATCC43255 wild-type (black spectra) or *ΔhadA* (gray spectra), after 48h of growth in MMM with 15mM [U-^13^C]threonine. Y-axes depict unitless NMR signal and X-axes depict ^13^C chemical shift in ppm for carbons 1-4 in butyrate (panels **A-D**) or 2-hydroxybutyrate (panels (**E-H**). Butyrate and 2-hydroxybutyrate isoptomers are shown in the upper right-hand corner of each panel with the carbon under analysis indicated by red shading. Dark and light blue arrows in the chemical structures indicate J-coupling signals from adjacent ^13^C carbons, and as reflected by the dotted lines over the spectra showing the associated split peaks. Labeled gray arrows indicate signals from non-target compounds. Splitting patterns at positions (**A**) C1, (**B**) C2, (**C**) C3, and (**D**) C4 indicate formation of [2,3,4-^13^C]butyrate and [U-^13^C]butyrate species in the wild-type condition only. (**E-H**) ^13^C-NMR signal of 2-hydroxybutyrate carbons un supernatants of *C. difficile* ATCC43255 wild-type (black spectra) or *ΔhadA* (gray spectra) grown with 15mM [U-^13^C]threonine. Splitting patterns at positions (**E**) C1, (**F**) C2, (**G**) C3, and (**H**) C4 reveal [2,3,4-^13^C]2-hydroxybutyrate and [U-^13^C]2-hydroxybutyrate species in similar proportions to butyrate.

### Deletion of the *hadA* 2-hydroxyisocaproate CoA transferase eliminates threonine’s reduction to butyrate

To confirm the reductive leucine pathway’s role in reductive threonine fermentation, we evaluated metabolism in a *ΔhadA* mutant. [U-^13^C]threonine metabolized by the *ΔhadA* mutant yielded a 5.65-fold increase in signal from the upstream substrate [^13^C]2-hydroxybutyrate (Fig. 3e-h, gray spectra), but no detectable [^13^C]butyrate (Fig. 3a-d, gray spectra).

### *P. bifermentans* threonine consumption *in vivo* further reduces dual redox substrates for *C. difficile’s* metabolism

*In vivo* studies by Girinathan, et al. demonstrated how the Stickland fermenter *Paraclostridium bifermentans* out-competes *C. difficile* for preferred amino acid substrates, including leucine and proline, rescuing a susceptible host from lethal disease[22, 23]. To examine *P. bifermentans’* effects on *C. difficile’s* threonine metabolism, we evaluated cecal levels of natural abundance threonine, 2-oxobutyrate, 2-aminobutyrate, and proprionate in gnotobiotic-and mice mono-colonized with *C. difficile, P. bifermentans*, or both species[9]. *C. difficile* depleted 69.6% of threonine in monocolonized mice (Fig. 4A), increasing levels of 2-oxobutyrate 77-fold (Fig. 4B) and 2-aminobutyrate 19-fold (Fig 4C). In contrast, *P. bifermentans* depleted 68.2% of threonine (Fig. 4A) and produced abundant propionate (Fig. 4D), a finding consistent with its known oxidative threonine metabolism to propionate, and absent reductive Stickland metabolism of the amino acid [21]. While threonine depletion remained comparable in the co-colonized state, 2-oxobutyrate and 2-aminobutyrate production by *C. difficile* fell by by 75% (Fig. 4C), in line with *P. bifermentans’* upstream consumption of threonine. In co-colonized mice, *P. bifermentans* effects repressed *C. difficile’s* threonine dehydratase expression by 52% at 20 hours of infection, and further by 43% at 24 hours (Fig. 4E), while threonine aldoase expression was not significantly altered (Fig. 4F). These findings extend *P. bifermentans’* metabolic competition to include repression of *C. difficile’s* theronine metabolism at the level of its gateway enzyme, threonine dehydratase.

**Figure 4.**
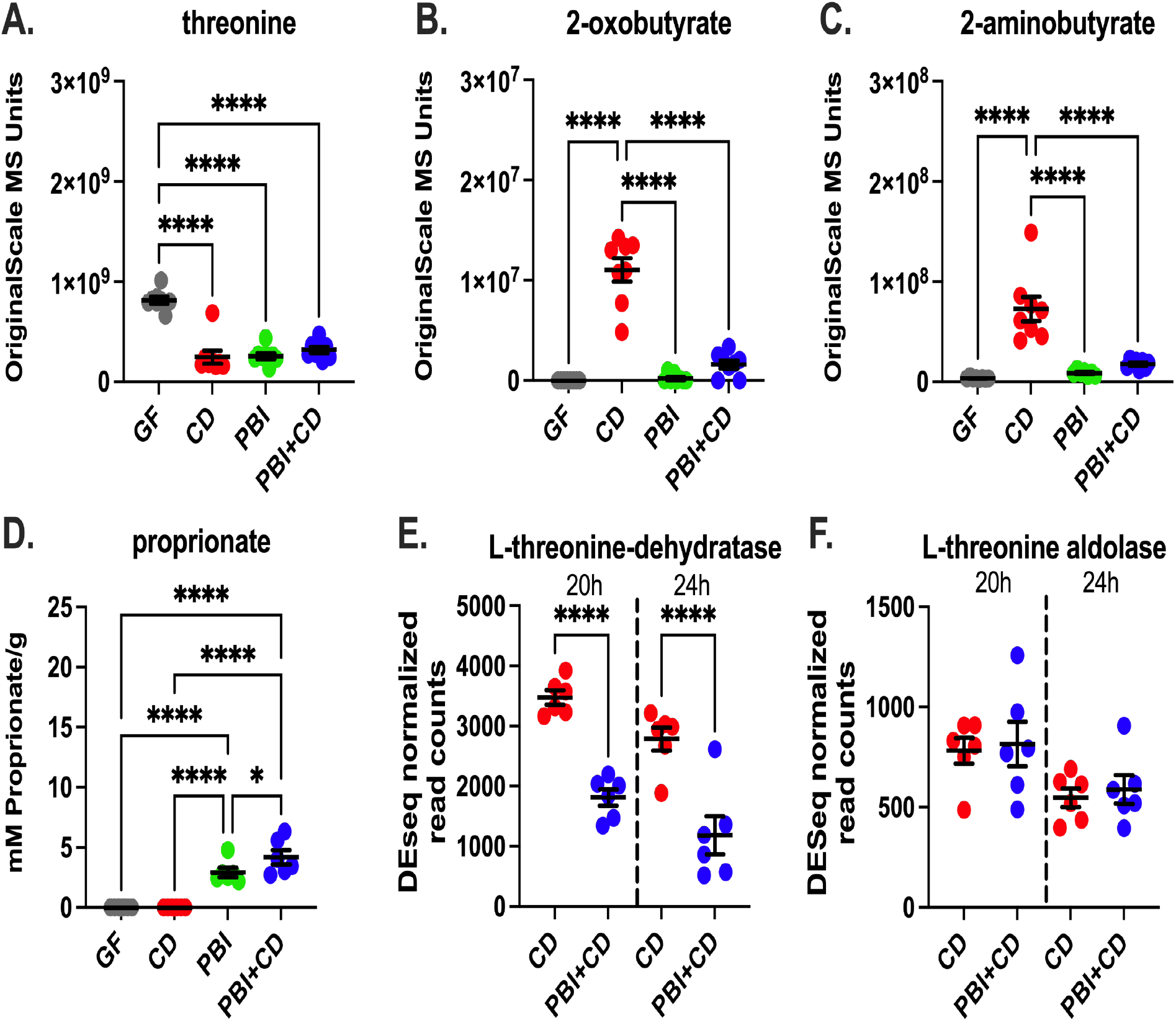
*P. bifermentans* represses *C. difficile’s* threonine metabolism *in vivo*. Analyses of metabolomic and metatranscriptomic data from Girinathan, et al. [22]. Panels A-C show cecal metabolomic data obtained by untargeted mass spectrometry in germfree mice (grey circles), mice mono-colonized with *P. bifermentans* for 7 days (green circles), *C. difficile* infected for 24h (blue circles) or co-colonzied mice (blue) after 7 days of PBI mono-colonization and 24h of *C. difficile* infection, for (A) threonine, (B) 2-oxobutyrate, (C) 2-aminobutyrate. Y-axis shows original scale mass spectrometry units, and conditions along the X-axis. Priopionate levels in panel D were measured by quantitative flame ionization detection (FID) for volatile short chain fatty acids to quantitate mM of propionate per grams of cecal contents (Y-axis); ****p<0.001. Panels (E) and show DESeq2 normalized read counts for *C. difficile’s* threonine-specific metabolic enzymes. Data shown at 20 and 24h of infection. (E) threonine dehydratase; DESeq2 adjusted p value at 20h betwen CD and PBI+CD = 0.0350 and at 24h = 0.0011, and (F) threonine aldolase, which showed no significant differences in expression levels among groups or timepoints.

## Discussion

We identify novel dynamics for Stickland threonine metabolism in *C. difficile*. Real-time whole-cell HRMAS ^13^C NMR with genome-scale metabolic modeling deconvoluted threonine’s dynamic support for changing cellular needs in nitrogen handling, energy production, redox balance, and growth. *C. difficile’s* initial deamination of threonine to 2-oxobutyrate releases ammonia in a redox-independent mechanism, conserving electron carriers for oxidative fermentations. Subsequent transamination of 2-oxobutyrate with amino groups donated from other amino acids produces 2-aminobutyrate, facilitating a parallel Stickland fermentation of the amino acid donor (Fig. 1f, Fig. 2a). Formation and depletion of 2-aminobutyrate also suggests a time-dependent role for threonine in cellular nitrogen cycling. Model solutions predicted 2-oxobutyrate’s transamination to 2-aminobutyrate in early metabolism, prior to 11 hours, providing an amino nitrogen sink to support deamination of other Stickland substrates, a role previously demonstrated for 3-carbon analogs in pyruvate’s transamination to alanine[20]. When coupled with the transamination of other Stickland amino acids, threonine’s conversion to 2-aminobutyrate provides an NAD-independent means for ammonia release, illustrating metabolic strategies used by Stickland fermenters to increase the efficiency of their amino acid metabolism. After 11 hours, 2-aminobutyrate’s re-conversion to 2-oxobutyrate shows how *C. difficile* further harnesses threonine-origin metabolites in late-stage reductive metabolism, yielding 2-hydroxybutyrate and butyrate, in addition to *n-*propanol from oxidatively-produced propionate. At this point, model predictions inferred transfer of 2-aminobutyrate’s amino groups to glycolytically-produced pyruvate to produce alanine, via alanine transaminase, or release as free ammonia via glutamate dehydrogenase (Fig. 1c,f). In this manner 2-aminobutyrate serves three functions as an efficient nitrogen sink, energy store, and lastly as a non-canonical Stickland amino acid supporting reductive metabolism.

Oxidative Stickland fermentation of 2-oxobutyrate to propionate, with one molecule of ATP (Fig. 2b), peaked over 6-16h, between points of maximal flux of preferred Stickland substrates such as leucine and proline, and prior to maximal flux through glycolytic pathways. As needs for reductive metabolism shifted, propionate’s reduction to *n-*propanol recycled 2 NADH to NAD+ (Fig. 2c). This overall pathway provides oxidative then reductive support, generating ATP, followed by net consumption of one electron pair in propionate’s reduction to support oxidative flux in other pathways. These reactions differ from those for leucine fermentation, where oxidatively-produced isovalerate is a terminal metabolite and does not support further reductive metabolism to isoamyl alcohol[20].

In later phases of growth, *C. difficile* reduces threonine to butyrate via a reductive pathway comparable to that described in *C. propionicum*[19] (Fig. 2d). Real-time HRMAS NMR detected the intermediate [^13^C]2-hydroxybutyrate with similar isotope composition to pooled [^13^C]butyrate. The dominant reductive productions, [U-^13^C]butyrate and [2,3,4-^13^C]butyrate, suggest that threonine’s carbon backbone remains intact during reduction, or is modified only through carboxyl group exchange on propionate, during dynamic equilibrium of the oxidative Stickland pathway. If butyrate production had occurred using ^13^C_2_ units from this pathway, populations of [1,2-^13^C]butyrate and [3,4-^13^C]butyrate would be expected from independent mixture with natural abundance acetyl-CoA from glycolytic, glycine, and cysteine fermentations. However, the absence of [1,2-^13^C]butyrate and [3,4-^13^C]butyrate confirms that ^13^C_2_ units from threonine cleavage did not appreciably contribute to butyrate’s production in the given media condition. In addition, threonine aldolase activity was detected via the production of [^13^C]glycine and [^13^C]ethanol (Fig. 2e), confirming a pathway by which threonine metabolism connects into central carbon metabolism and biomass production.

We propose that *C. difficile’s* reductive leucine pathway performs functions of the lactate-acrylate-propionate systems in other *Clostridia* to support reductive threonine metabolism to butyrate. In-frame deletion *of C. difficile’s* 2-hydroxyisocaproate CoA transferase in the reductive leucine pathway completely abolished [^13^C]butyrate from [^13^C]threonine, accompanied by accumulation of the upstream intermediate [^13^C]2-hydroxybutyrate. Studies by Kim et al. demonstrated that butyryl-CoA is not accepted as the CoA donor in the 2-hydroxyisocaproate CoA transferase reaction[24]. Therefore, a different mechanism, such as via phosphobutyryl transferase and butyrate kinase, may facilitate butyrate’s release from CoA, with the 2-hydroxyisocaproate CoA transferase catalyzing the transfer of CoA to 2-hydroxybutyrate.

The reductive leucine pathway’s promiscuity enhances its contributions to *C. difficile’s* versatile metabolic toolbox, including to ferment prevalent amino acids in gut contents[11-13]. Prior induction of reductive leucine metabolism for the preferred substrate leucine would provide an already induced pathway to consume threonine, with gateway control of threonine flux via *C. difficile’s* threonine transport systems and threonine dehydratase to convert threonine to 2-oxobutyrate. Whereas previous studies examining leucine and proline fermentation identify Stickland fermentations as an early energy source[20], we demonstrate how reductive threonine fermentation supports growth and metabolism into transition and stationary phases of growth.

The 2-hydroxyisocaproyl-CoA dehydratase in the reductive leucine pathway has previously been identified as a key target of the antimicrobial metronidazole[25]. While metronidazole remains a non-preferred therapeutic due to its severe suppression of protective microbiota and risks for recurrent infections[26], our findings suggest that the reductive leucine pathway offers an additional metabolic target amenable to alternate therapies targeting *C. difficile’s* electron bifurcating enzymes.

We further demonstrate dynamics of *C. difficile’s* threonine fermentation *in vivo* illustrating primary conversion to 2-aminobutyrate during early colonization and infection. *P. bifermentans’* threonine metabolism to propionate reduces available pools of a preferred Stickland redox substrate for *C. difficile*, acting in combination within its broader metabolism to create a nutrient-depleted environment, limiting the pathogen’s ability to colonize and infect the gut.

*In vivo*, threonine’s fermentation to butyrate by commensal Stickland fermenters provides an alternate source of this host-active metabolite, and one that can arise independently of dietary carbohydrates and complex fibers administered to elevate levels [27, 28]. These pathways offer further consideration for metabolic interventions in pathogens carrying the *had* pathway, such as in *C. botulinum* and *C. sordellii*, and for solvogenic species used to generate butyrate and butanol from complex feedstocks, such as for carrying strains of *C. butyricum*.

Threonine’s complex dynamics also advanced approaches for conducting dFBA in obligate anaerobes under anoxic and reducing conditions, including support to model intermediate accrual and consumption, as occured with 2-aminobutyrate and propionate flux. We implemented additional enhancements to better model flux for metabolites with multiple metabolic origins, such as butyrate, which arises from multiple sources including glucose and threonine. These enhancements include methods to infer component trajectories across experiments using different [U-^13^C] inputs to constrain the flux of contributing reactions that produce common metabolites, whether they originate from distinct or converging pathways. These enhancements improved the accuracy of dFBA solutions in modeling threonine fermentation and further enhance the generalizability of our dFBA framework for anaerobe metabolism.

We resolve *C. difficile’s* dynamic metabolism threonine’s including to colonize host ecosystems. HRMAS NMR-informed dFBA solutions predicted coordinated recruitment of multiple threonine utilization pathways with systems regulating energy generation, redox balance, and nitrogen cycling. We show that threonine fermentation leverages defined gateway enzymes, in threonine dehydratase and threonine aldolase, to control downstream metabolic flux through promiscuous systems including oxidative Stickland enzymes, and the reductive leucine and butyrate fermentation pathways. These findings demonstrate how *C. difficile* leverages this underlying metabolic diversity to ferment gut-available nutrients that drive its colonization and progression to disease.

## Methods

### Strains

*C. difficile* ATCC 43255 ΔPaLoc [20] was used for time-series NMR analyses. For studies evaluating butyrate production in stationary phase supernatants, an in-frame deletion of the *hadA* strain in *C. difficile* ATCC 43255 was generated, and uniquness of the mutant confirmed by genome sequencing of the isogenic mutant strain. The strain was shown by gas chromatography with flame ionization detection to produce no isocaproate (data not shown).

### Strain culture conditions

*C. difficile* MMM was prepared as described in Pavao, et al [20]. Preparations with labeled threonine used 15 mM L-[U-^13^C]threonine (Sigma-Aldrich) in place of natural-abundance L-threonine. The elevated threonine concentration was selected to enable enhanced detection of minor metabolic products.

Tryptone-Yeast-Glucose (TYG) media was prepared according to the formulation in Pavao, et al. [20].

Prior to serial real-time whole-cell HRMAS-NMR experiments, cells were grown anaerobically for 16 hours in MMM. Cells were spun and washed three times in pre-reduced sterile PBS. The cell suspension was diluted to introduce 100,000 cells into an sealed insert for HRMAS Kel-F 4mm rotor (Bruker BioSpin Corporation) containing growth media with ^13^C-label substrates. The insert was sealed and removed from the anaerobic chamber for NMR analyses.

### HRMAS NMR and spectrum analyses

HRMAS NMR measurements were performed on a Bruker Avance III HD 600 MHz spectrometer (Bruker BioSpin Corporation). Acquisition was performed as described in Pavao, et al. [20].

Individual free induction decay (.fid) files were processed as described in [NatChemBio], a graphical user interface was added to the processing pipeline to enhance usability. All custom code is available on the GitHub repository at https://github.com/Massachusetts-Host-Microbiome-Center/nmr-cdiff.

### Metabolic modeling

dFBA was performed using an enhanced copy of icdf843 [20] containing the novel threonine-to-butyrate fermentation pathway.

Exchange fluxes were estimated according to the procedure in Pavao, et al. [20]. To support more accurate modeling of threonine fermentation, we made the following enhancements to the codebase:

1. Added support for non-logistic curves. In cases of intermediate accrual and consumption, as used to model 2-aminobutyrate and propionate flux, a sum of two logistic curves was used.
2. To improve modeling of metabolites with multiple metabolic origins, support was added for a sum of trajectories across separate experiments with different [U-^13^C] input substrates to constrain the flux of common metabolites generated by distinct or converging pathways. This feature was used to support acetate and butyrate trajectories.
3. To better constrain the converging pathways in this scenario, we have also included the option to define flux through contributing reactions that support the formation of a particular product. This feature was used to assess feasibility of butyrate-forming fluxes through the reductive leucine pathway and the classical acetyl-CoA pathway.

Simulations used exchange flux constraints from Pavao, et al. [20] in addition to those calculated from the [U-^13^C]threonine HRMAS NMR experiments performed in this study. Standard curves to estimate concentration from HRMAS NMR integrated signal used [U-^13^C]butyrate, [U-^13^C]propionate, and [U-^13^C]1-propanol (Cambridge Isotope Laboratories, Inc.). Ethyl [U-^13^C]2-hydroxybutyrate (Cambridge Isotope Laboratories, Inc.) approximated 2-hydroxybutyrate and 2-aminobutyrate signals, as neither compound was commercially available as a U-^13^C preparation.

dFBA was otherwise performed as described in Pavao, et al. [20] using the COBRApy toolbox and custom Python scripts.

### Theronine metabolite and *C. difficile* metabolic gene analyses in defined-association mice

The untargeted metabolomic datasets from Girinathan, et al. were used to evaluate levels of threonine, 2-oxobutyrate (labeled as alpha-ketobutyrate in the original dataset), 2-aminobutyrate, and propionate, in gnotobiotic mice, animals monocolonized for 7 days with *P. bifermentans*, 24 hours of mono-colonization with *C. difficile*, or 7 days with *P. bifermentans*, followed by 24h of infection with *C. difficile* (co-colonized). Datapoints were plotted in Prism and significance among groups evaluated by ANOVA with Holm-Sidak multi-hypothesis correction. A p value <0.05 was considered significant. Analyses of *C. difficile’s* threonine dehydratase and threonine aldolase evaluated the DESeq2 normalized read counts among the above groups. Datapoints were plotted in Prism and evaluated by the DESeq2 adjusted p values among pairwise comparisons.

## Achnowledgements

We thank Mary Delaney for conducting short chain fatty acid analyses of the wild-type and Δ*hadA* mutant strains. This work was supported by grants R03AI174158, R01AI153605, and P30DK34854 (LB), OD023406, AG070257, and CA273010 (LLC); and from the Massachusetts Life Sciences Center (LB, LLC), and MGH A. A. Martinos Center for Biomedical Imaging (LLC, LB)

## Notes

### Competing Interest Statement

The authors have declared no competing interest.

## References

1. Wells, C.L. and T.D. Wilkins, Clostridia: Sporeforming Anaerobic Bacilli, in Medical Microbiology, S. Baron, Editor. 1996: Galveston (TX).

2. Morrison, D.J. and T. Preston, Formation of short chain fatty acids by the gut microbiota and their impact on human metabolism. Gut Microbes, 2016. 7(3): p. 189–200.

3. Lee, S.Y., et al., Fermentative butanol production by Clostridia. Biotechnol Bioeng, 2008. 101(2): p. 209–28.

4. Neumann-Schaal, M., D. Jahn, and K. Schmidt-Hohagen, Metabolism the Difficile Way: The Key to the Success of the Pathogen Clostridioides difficile. Front Microbiol, 2019. 10: p. 219.

5. Gencic, S. and D.A. Grahame, Diverse Energy-Conserving Pathways in Clostridium difficile: Growth in the Absence of Amino Acid Stickland Acceptors and the Role of the Wood-Ljungdahl Pathway. J Bacteriol, 2020. 202(20).

6. Kazamias, M.T. and J.F. Sperry, Enhanced fermentation of mannitol and release of cytotoxin by Clostridium difficile in alkaline culture media. Appl Environ Microbiol, 1995. 61(6): p. 2425–7.

7. Nakamura, S., et al., Carbohydrate fermentation by Clostridium difficile. Microbiol Immunol, 1982. 26(2): p. 107–11.

8. Pavao, A., et al., Reconsidering the in vivo functions of Clostridial Stickland amino acid fermentations. Anaerobe, 2022:p. 102600.

9. Girinathan, B.P., et al., In vivo commensal control of Clostridioides difficile virulence. Cell Host & Microbe, 2021. 29(11): p. 1693–1708.e7.

10. Arrieta-Ortiz, M.L., et al., Predictive regulatory and metabolic network models for systems analysis of Clostridioides difficile. Cell Host & Microbe, 2021. 29(11): p. 1709–1723.e5.

11. Moughan, P.J. and S.M. Rutherfurd, Gut luminal endogenous protein: implications for the determination of ileal amino acid digestibility in humans. Br J Nutr, 2012. 108 Suppl 2: p. S258–63.

12. Gum, J.R., Jr., et al., The human MUC2 intestinal mucin has cysteine-rich subdomains located both upstream and downstream of its central repetitive region. J Biol Chem, 1992. 267(30): p. 21375–83.

13. Puiman, P.J., et al., Intestinal threonine utilization for protein and mucin synthesis is decreased in formula-fed preterm pigs. J Nutr, 2011. 141(7): p. 1306–11.

14. Mead, G.C., The amino acid-fermenting clostridia. J Gen Microbiol, 1971. 67(1): p. 47–56.

15. Aguirre, A.M., et al., Bile acid-independent protection against Clostridioides difficile infection. PLoS Pathog, 2021. 17(10): p. e1010015.

16. Neumann-Schaal, M., et al., Time-resolved amino acid uptake of Clostridium difficile 630Δerm and concomitant fermentation product and toxin formation. BMC Microbiol, 2015. 15: p. 281.

17. Barker, H.A. and T. Wiken, The origin of butyric acid in the fermentation of threonine by Clostridium propionicum. Arch Biochem, 1948. 17(1): p. 149–51.

18. Barker, H.A., CHAPTER 3 - Fermentations of Nitrogenous Organic Compounds, in Metabolism, I.C. Gunsalus and R.Y. Stanier, Editors. 1961, Academic Press. p. 151–207.

19. Hofmeister, A.E. and W. Buckel, (R)-lactyl-CoA dehydratase from Clostridium propionicum. Stereochemistry of the dehydration of (R)-2-hydroxybutyryl-CoA to crotonyl-CoA. Eur J Biochem, 1992. 206(2): p. 547–52.

20. Pavao, A., et al., Elucidating dynamic anaerobe metabolism with HRMAS (13)C NMR and genome-scale modeling. Nat Chem Biol, 2023. 19(5): p. 556–564.

21. Elsden, S.R. and M.G. Hilton, Volatile acid production from threonine, valine, leucine and isoleucine by clostridia. Arch Microbiol, 1978. 117(2): p. 165–72.

22. Girinathan, B.P., et al., In vivo commensal control of Clostridioides difficile virulence. Cell Host Microbe, 2021.

23. Arrieta-Ortiz, M.L., et al., Predictive regulatory and metabolic network models for systems analysis of Clostridioides difficile. Cell Host Microbe, 2021. 29(11): p. 1709–1723 e5.

24. Kim, J., et al., Characterization of (R)-2-hydroxyisocaproate dehydrogenase and a family III coenzyme A transferase involved in reduction of L-leucine to isocaproate by Clostridium difficile. Appl Environ Microbiol, 2006. 72(9): p. 6062–9.

25. Kim, J., D. Darley, and W. Buckel, 2-Hydroxyisocaproyl-CoA dehydratase and its activator from Clostridium difficile. FEBS J, 2005. 272(2): p. 550–61.

26. Allegretti, J.R., et al., Clinical Predictors of Recurrence After Primary Clostridioides difficile Infection: A Prospective Cohort Study. Dig Dis Sci, 2020. 65(6): p. 1761–1766.

27. O’Keefe, S.J., Tube feeding, the microbiota, and Clostridium difficile infection. World J Gastroenterol, 2010. 16(2): p. 139–42.

28. Mortensen, P.B., et al., Colonic fermentation of ispaghula, wheat bran, glucose, and albumin to short-chain fatty acids and ammonia evaluated in vitro in 50 subjects. JPEN J Parenter Enteral Nutr, 1992. 16(5): p. 433–9.

